# APE1 active site residue Asn174 stabilizes the AP-site and is essential for catalysis

**DOI:** 10.1101/2025.01.14.633034

**Authors:** Kaitlin M. DeHart, Nicole M. Hoitsma, Spencer H. Thompson, Veniamin A. Borin, Pratul K. Agarwal, Bret D. Freudenthal

## Abstract

Apurinic/Apyrimidinic (AP)-sites are common and highly mutagenic DNA lesions that can arise spontaneously or as intermediates during Base Excision Repair (BER). The enzyme apurinic/apyrimidinic endonuclease 1 (APE1) initiates repair of AP-sites by cleaving the DNA backbone at the AP-site via its endonuclease activity. Here, we investigated the functional role of the APE1 active site residue N174 that contacts the AP-site during catalysis. We analyzed the effects of three rationally designed APE1 mutations that alter the hydrogen bonding potential, size, and charge of N174: N174A, N174D, and N174Q. We found impaired catalysis of the APE1_N174A_ and APE1_N174D_ mutants due to disruption of hydrogen bonding and electrostatic interactions between residue 174 and the AP-site. In comparison, the APE1_N174Q_ mutant was less impaired due to retaining similar hydrogen bonding and electrostatic characteristics as N174 in wild-type APE1. Structures and computational simulations further revealed that the AP-site was destabilized within the active sites of the APE1_N174A_ and APE1_N174D_ mutants due to loss of hydrogen bonding between residue 174 and the AP-site. Cumulatively, we show that N174 stabilizes the AP-site within the APE1 active site through hydrogen bonding and electrostatic interactions to enable effective catalysis. These findings highlight the importance of N174 in APE1’s function and provide new insights into the molecular mechanism by which APE1 processes AP-sites during DNA repair.

## Introduction

Oxidative DNA damage can result from exposure to oxidizing agents, either in the environment or generated as byproducts of cellular metabolism (1–4). Repair of oxidative DNA damage is essential to protect genome stability and prevent deleterious mutations that give rise to multiple human diseases (5–7). This repair is primarily accomplished via the Base Excision Repair (BER) pathway (8–11). During BER, damaged DNA nucleobases are identified and removed by a damage-specific DNA glycosylase that generates baseless sugar moieties known as apurinic/apyrimidinic (AP)-sites (12–15). AP-sites are then processed by Apurinic/Apyrimidinic Endonuclease 1 (APE1), which cleaves the phosphodiester DNA backbone at the 5’ end of the AP-site using its endonuclease activity (16–20). This substrate is subsequently processed by downstream BER proteins to complete repair of the damaged base (11,21,22). APE1 cleavage of AP-sites is an essential process to protecting genome stability during DNA repair as the knockout of APE1 is embryonic lethal (18). Similarly, expression of inactive APE1 variants sensitizes cells to DNA damaging agents (23–26). Because APE1 cleavage of AP-sites constitutes an important step in DNA repair, investigation of the APE1 cleavage mechanism and key active site residues is of interest to more broadly understand how DNA damage is processed by this essential enzyme.

Previous high resolution x-ray crystal structures of APE1 bound to AP-site containing DNA, both before and after cleavage, have provided insight into the mechanism of APE1 cleavage and APE1 active site organization (Fig. 1A,B) (27–29). Upon binding an AP-site, APE1 flips the AP-site out of the DNA double helix and into an extrahelical position within the compact APE1 active site (Fig. 1A) (27,28). The AP-site is positioned near active site residues E96, Y171, D308, and H309 along with a single Mg^2+^ ion (Fig. 1B) (30,31). This places the phosphate backbone of the AP-site in position for cleavage through a one metal hydrolysis reaction (28,31,32). To coordinate this reaction, a water molecule is positioned for inline nucleophilic attack on the phosphorus atom of the AP-site through hydrogen bonding interactions with residues N212 and D210 (Fig. 1B) (28,33). D210 deprotonates the water in this position, activating the water for nucleophilic attack and initiating hydrolysis (28,29). After cleavage, the DNA product consists of a 5’-sugar phosphate terminus and a 3’-hydroxyl terminus. In this product state, N212 and D210 are within hydrogen bonding distance of the 5’ phosphate terminus of the cleaved AP-site (Fig. 1C) (28). Similarly, E96, Y171, D308, H309 and the Mg^2+^ cofactor maintain coordination of the phosphate group, now the 5’ terminus, as well as the newly formed 3’ hydroxyl terminus (Fig. 1C) (27,28,30).

**Figure 1.**
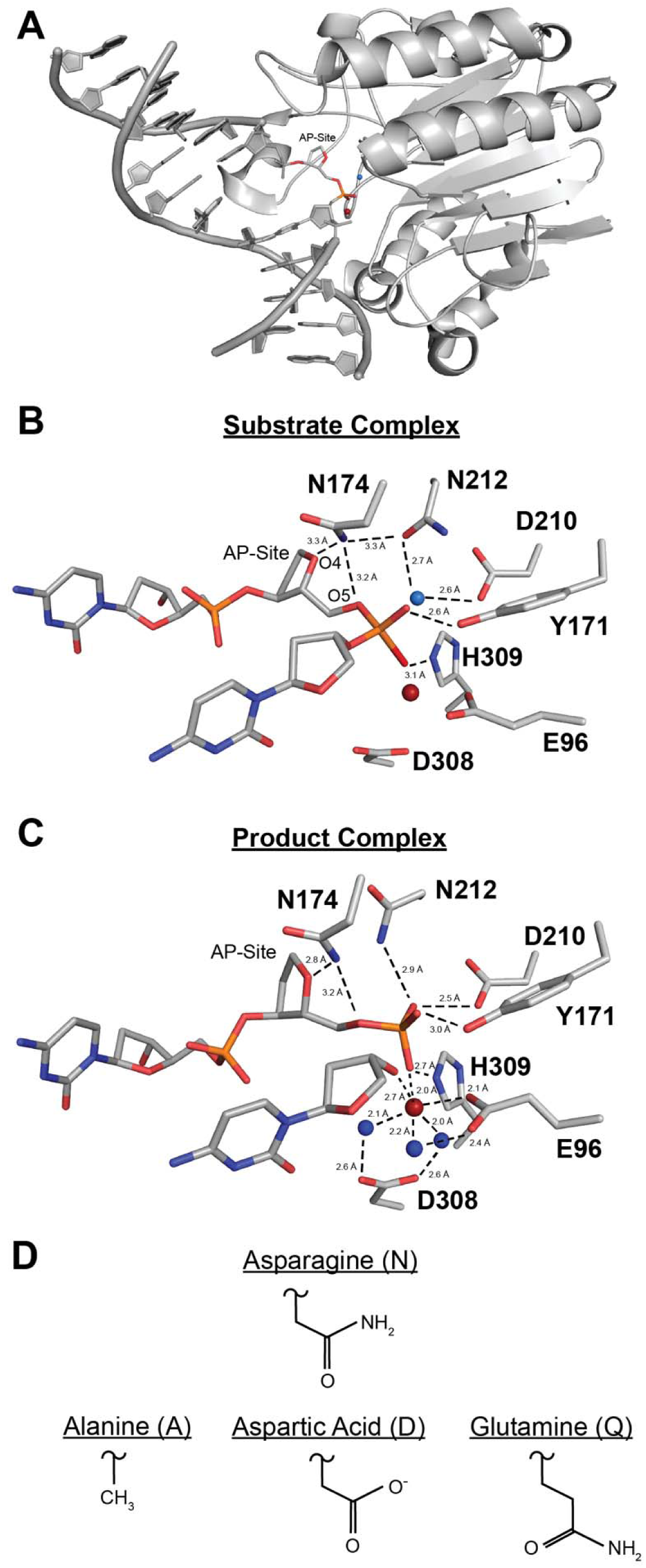
The structure of APE1_WT_ and N174 mutations. (A) Overview of APE1_WT_ bound to a DNA substrate containing an AP-site (PDB ID: 5DFI). (B) Close up of the APE1 active site substrate complex (PDB ID: 5DFI) bound to a DNA substrate containing an AP-site. Waters shown as blue spheres and a modeled Mg^2+^ ion is shown as a red sphere. 5DFI lacks a metal in the structure, but Mg^2+^ ions coordinate the APE1_WT_ active site in both substrate and product complexes. We have estimated the position of Mg^2+^ in the substrate complex by alignment with its position in the product complex 5DFF (RMSD=0.301 of 3945 atoms). (C) Close up of the active site of the APE1_WT_ product complex (PDB ID: 5DFF). Water shown as blue spheres, Mg^2+^ shown as a red sphere. (D) Side chains of mutations at residue 174: N174A, N174D, and N174Q.

While many of the residues in the APE1 active site have been investigated, there is a glaring lack of knowledge about active site residue N174. X-ray crystal structures of APE1 indicate that N174 contacts the AP-site (Fig. 1B, C), but the role of N174 during AP-site cleavage is unknown. Interestingly, N174 is highly conserved among chordate APE1 and shares similarity with Q112 in the APE1 homolog and primary AP-site endonuclease in *E. coli*, EXOIII (Fig. S1) (34–40). This suggests N174 may be required for APE1 cleavage of AP-sites. In the human APE1 substrate complex, N174 is within hydrogen bonding distance of N212 and the AP-site bridging oxygen (O5) and deoxyribose sugar oxygen (O4) (Fig. 1B, Fig S2A, S2B) (27,28). In the product complex, N174 maintains hydrogen bonding distance to O4 and O5 of the AP-site after cleavage (Fig. 1C) (27,28). Because of its unique position near the AP-site, as well as its direct interaction with the phosphodiester backbone at this location, we hypothesized that N174 may stabilize substrate DNA during the cleavage reaction. To assess the role of N174 during APE1 catalysis, we rationally designed three mutations of N174: N174A, N174D, and N174Q. We investigated these APE1 variants using kinetic, structural and computational approaches. From these studies we demonstrated that N174 is required to efficiently position the AP-site in the APE1 active site during catalysis. In addition, N174 establishes an electrostatic environment required for catalysis. These observations expound upon the mechanism of the APE1 cleavage reaction and provide insight into the role of N174 during AP-site cleavage by APE1.

## Results

To determine the role of N174 during APE1 cleavage, we generated three APE1 mutants (N174A, N174D, and N174Q) that altered distinct attributes of the amino acid side chain at residue 174: hydrogen bonding potential, charge, and size (Fig. 1D). The N174A mutation has no hydrogen bonding potential, no charge or polarity, and has a smaller side chain length than N174. The N174D mutation has no hydrogen bonding potential and has a negative charge compared to N174 but is the same size as N174. Finally, the N174Q mutation has a similar hydrogen bonding potential and charge compared to N174 but has a larger side chain size than N174. Using these three APE1 mutants, we analyzed how hydrogen bonding, charge, and size of N174 affect APE1 cleavage of AP-sites.

### N174A and N174D Mutations Inhibit APE1 Cleavage of AP-sites

Prior to cleavage, N174 is within hydrogen bonding distance of O4 and O5 of the AP-site (Fig. 1B). We hypothesized that mutation of this residue would destabilize the substrate DNA and impair catalysis. To test this, we utilized multi-turnover, pre-steady state kinetic analysis to determine whether mutating N174 affected the rates of APE1 cleavage (*k_obs_*) and product release (*k_ss_*) (Table 1, Fig. S2). Under multi-turnover conditions, APE1 demonstrates biphasic kinetics for product formation (Fig. S2A). The initial burst phase of product formation corresponds to the APE1 cleavage rate constant (*k*_obs_) while the second phase represents the steady state rate constant (*k_ss_*) corresponding to product release (41).

**Table 1.**
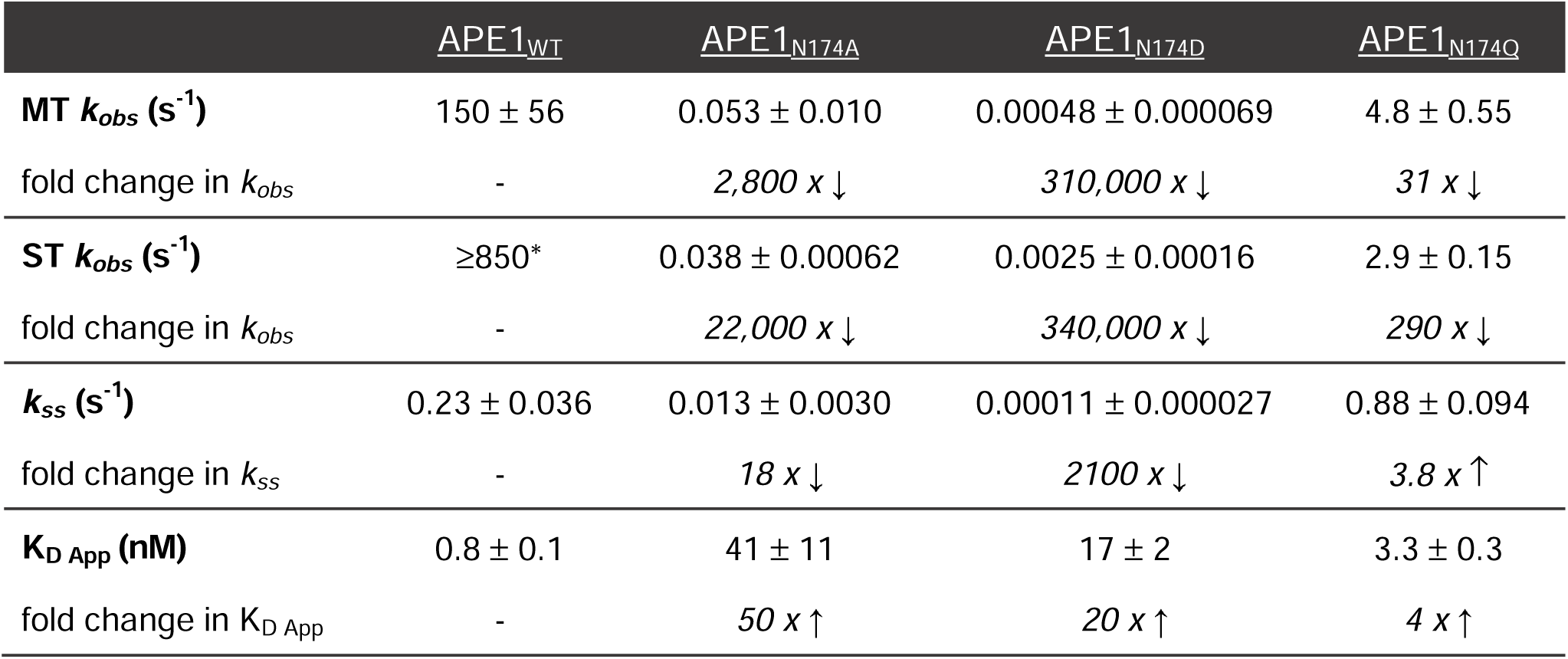
Summary of kinetic rate constants and binding affinities for APE1_WT_, APE1_N174A_, APE1_N174D_, and APE1_N174Q_ Rate constants (mean ± SE, N=3) corresponding to cleavage (*k_obs_*) and product release (*k_ss_)* were determined by multiple turnover (MT) and/or single turnover (ST) kinetic analysis. Apparent binding affinities determined by EMSA. Fold changes are relative to APE1_WT_ values. * As determined by the following reference: (41). See methodology for additional information.

Analysis of the burst phase of multiple turnover kinetic analysis revealed that each mutant exhibited a reduced *k*_obs_ compared to APE1_Wild Type (WT)_ (Table 1, Fig. S2). Under multi-turnover conditions, the *k_obs_* for APE1_WT_ was 150 ± 56 s^−1^, consistent with previously published findings (41). The *k_obs_*for the APE1_N174A_ mutant was 0.053 ± 0.010 s^−1^, corresponding to a 2,800-fold decrease in the *k_obs_*compared to APE1_WT_. The *k_obs_*of the APE1_N174D_ mutant was 0.00048 ± 0.000069 s^−1^, corresponding to a 310,000-fold decrease compared to APE1_WT_. Finally, the *k_obs_* of the APE1_N174Q_ mutant was 4.8 ± 0.55 s^−1^, corresponding to a 31-fold decrease compared to APE1_WT_. Thus, N174A and N174D mutations resulted in a drastically reduced *k_obs_* and the N174Q mutation resulted in a moderately reduced *k_obs_*. This finding indicates that N174 is necessary for APE1 cleavage.

The *k_obs_*values obtained for all APE1 mutants from multi-turnover experiments were validated by performing single-turnover kinetic analysis (Table 1, Fig. S3). Under single turnover conditions, the APE1_N174A_ mutant had a *k_obs_* of 0.038 ± 0.00062 s^−1^, corresponding to a 22,000-fold reduction in *k_obs_*compared to APE1_WT_ which is estimated to be ≥850 s^−1^ (41). The APE1_N174D_ mutant had a *k_obs_*of 0.0025 ± 0.00016 s^−1^, corresponding to a 340,000-fold reduction in *k_obs_* compared to APE1_WT_. Lastly, the APE1_N174Q_ mutant had a *k_obs_* of 2.9 ± 0.15 s^−1^, corresponding to a 290-fold reduction in *k_obs_* compared to APE1_WT_. Importantly, the *k_obs_* values obtained for all APE1 mutants determined by single-turnover kinetic analysis were consistent with values determined by multi-turnover kinetic analysis. Cumulatively, the multi-turnover and single-turnover kinetics indicate that residue N174 is key for APE1 cleavage.

Product release has been determined to be the rate-limiting step of APE1 catalysis, limiting enzyme turnover, and catalytic cycling (41). To determine the contribution of N174 to the product release step of APE1 catalysis, we assessed the steady state phase of our multi-turnover kinetics for all N174 APE1 mutants (Table 1, Fig. S2). APE1_WT_ had a *k_ss_* of 0.23 ± 0.036 s^−1^. The *k_ss_* of the APE1_N174A_ mutant was 0.013 s^−1^ ± 0.0030, corresponding to an 18-fold reduction compared to APE1_WT_. The *k_ss_*of the APE1_N174D_ mutant was 0.00011 ± 0.000027 s^−1^, corresponding to a 2,100-fold reduction compared to APE1_WT_. In contrast, the *k_ss_* of the APE1_N174Q_ mutant was 0.88 s^−1^ ± 0.094, corresponding to a 4-fold increase compared to APE1_WT_. Together, our kinetic analysis indicates that the N174A and N174D mutations impaired both the cleavage and product release rates of APE1. This indicates that the hydrogen bonding potential and charge that is altered in these mutants is affecting APE1 cleavage. In contrast, N174Q mutation which maintains the hydrogen bonding and charge, more moderately impacted the cleavage and product release rates of APE1. These findings are consistent with evidence that the hydrogen bonding and charge of N174 are required for catalysis in WT APE1.

To determine if N174 contributes to the ability of APE1 to bind DNA, we performed electrophoretic mobility shift assays (EMSAs). These experiments enabled us to determine the apparent binding affinity (*K*_D App_) of each APE1 mutant for DNA containing an AP-site (Table 1, Fig. S4). Determination of the *K*_D App_ would also allow us to determine if the reductions in *k_obs_* or *k_ss_* observed in the N174 APE1 mutants could result from a reduction in binding affinity for AP DNA. APE1_WT_ had a *K_D App_* of 0.8 ± 0.1 nM. The APE1_N174A_ mutant had a *K*_D App_ of 41 ± 11 nM, corresponding to a 50-fold decrease in apparent binding affinity compared to APE1_WT_. The APE1_N174D_ mutant had a *K*_D App_ of 17 ± 2 nM, corresponding to a 20-fold decrease in apparent binding affinity compared to APE1_WT_. And lastly, the APE1_N174Q_ mutant had a *K*_D App_ of 3.3 ± 0.3 nM, corresponding to a 4-fold decrease in apparent binding affinity compared to APE1_WT_. This indicates that residue N174 does contribute to AP DNA binding, though the effect on the *K*_D App_ is modest compared to what we observed for *k_obs_*and *k_ss_*.

### N174A Mutation Destabilizes the AP-site within the APE1_N174A_ Active Site

Previously solved x-ray crystal structures of the APE1_WT_ active site revealed that N174 is within hydrogen bonding distance of O4 and O5 of the AP-site prior to cleavage (Fig. 1B) (27,28). Therefore, we predicted that N174 would contribute to APE1 cleavage by stabilizing substrate DNA during catalysis. To test this, we crystallized the APE1_N174A_ mutant protein bound to a 21-mer double-stranded DNA oligo containing a centrally located AP-site analog tetrahydrofuran (THF). This substrate complex diffracted to 2.29 Å resolution in the P1 space group (Fig. 2A-B, Table 2). We have clear density for the N174A mutation as well as the AP-site, which is positioned within the APE1_N174A_ mutant active site (Fig. 2A-B). Overlay of the APE1_WT_ and APE1_N174_ substrate complexes revealed similar active sites, with no notable differences in the position of key active site residues (RMSD=0.236 of 1767 atoms, Fig. 2C) (28). In contrast, shifts to the DNA within the active site are observed, with the AP-site, phosphate backbone, and surrounding DNA adopting a different position in the APE1_N174A_ mutant compared to APE1_WT_ (Fig. 2D). Specifically, the AP-site phosphate group is shifted 1.1 Å out of the APE1 active site in the APE1_N174A_ mutant compared to APE1_WT_ (Fig. 2D). This shift increases the distance between the phosphate of the AP-site and the nucleophilic water, ultimately resulting in misalignment of the phosphate backbone for attack by the nucleophilic water (Fig. 2E). This loss of alignment does not appear to arise from changes in position of active site residues since the positions of both the nucleophilic water and the active site residues which coordinate it, N212 and D210, are not changed in the APE1_N174A_ mutant (Fig. 2C). Based on these results, we conclude that the N174A mutation compromises hydrogen bonding between residue 174 and O4 and O5 of the AP-site. This allows the DNA to shift away from the active site in the APE1_N174A_ mutant. Therefore, in APE1_WT_, the hydrogen bonds between N174 and the AP-site are necessary to position the phosphate backbone of the AP-site within the APE1 active site to support nucleophilic attack during catalysis.

**Figure 2.**
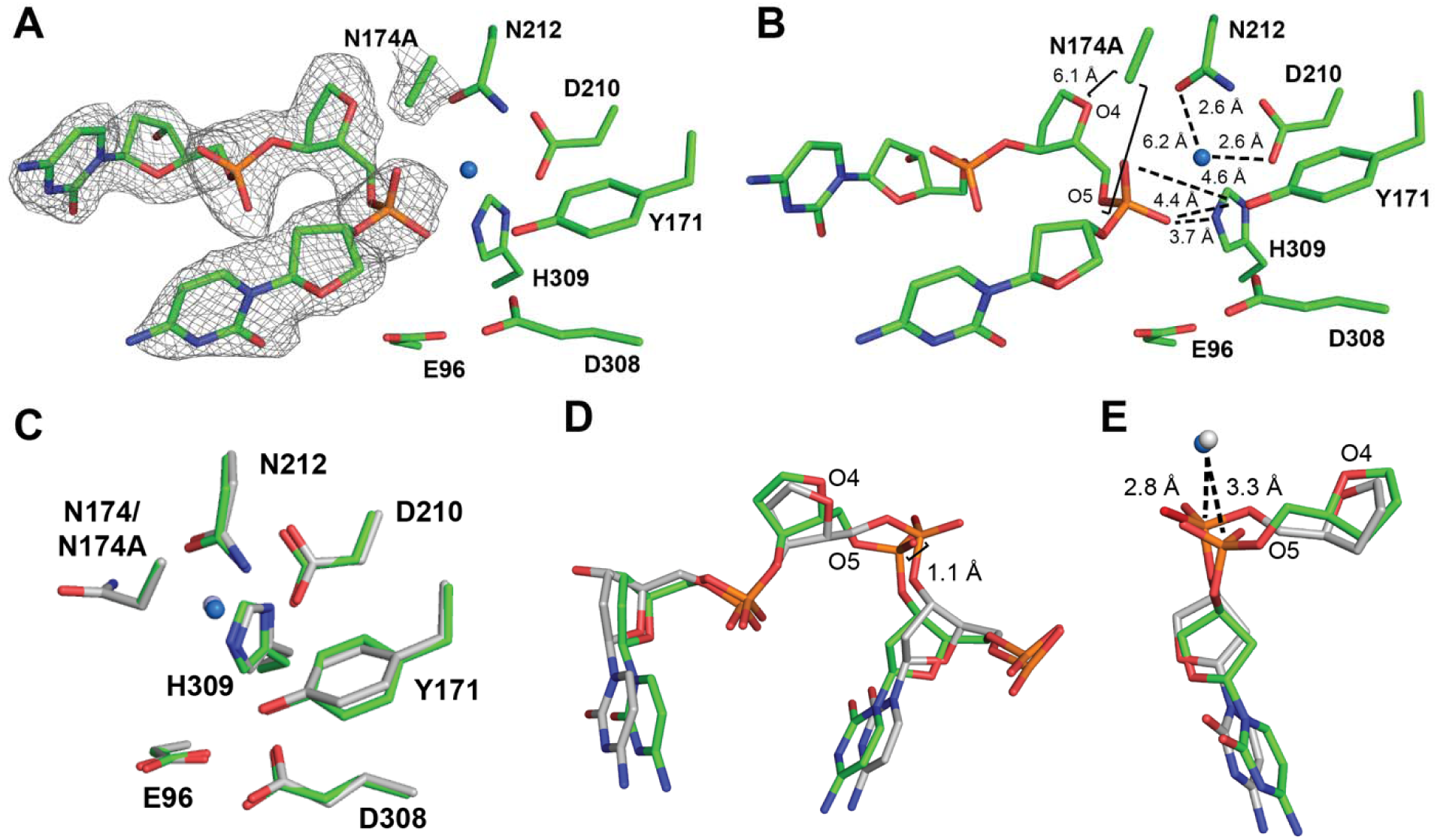
The APE1_N174A_ mutant substrate complex. (A-B) Active site of the APE1_N174A_ substrate complex crystal structure shown in green. Density is shown as grey mesh. Waters are shown as blue spheres. (C) Active site residues of the APE1_N174A_ substrate complex crystal structure compared to active site residues of the APE1_WT_ substrate complex shown in gray (5DFI). (D) The DNA backbone in the APE1_N174A_ mutant compared to APE1_WT_. (E) Distance between the nucleophilic water and DNA backbone increases in the APE1_N174A_ substrate complex crystal structure shown as a blue sphere compared to APE1_WT_ shown as a grey sphere.

**Table 2.** X-ray Structure Data Collection and Atomic Model Refinement Statistics Parentheses indicate value for highest resolution shell.

|  | **APE1<sub>N174A</sub>:**<br>**Substrate DNA**<br>**Complex** | **APE1<sub>N174A</sub>:**<br>**Product DNA**<br>**Complex** | **APE1<sub>N174D</sub>:**<br>**Product DNA**<br>**Complex** | **APE1<sub>N174D</sub>:**<br>**Product DNA**<br>**Complex** |
| --- | --- | --- | --- | --- |
| **Space group** | P1 | P1 | P1 | P43 |
| Cell dimensions |  |  |  |  |
| *a*, *b*, *c* (Å) | 43.9, 60.5, 73.1 | 44.3, 61.8, 72.6 | 44.2, 60.8, 73.6 | 153.4, 153.4, 45.4 |
| $\alpha$, $\beta$, $\gamma$ (°) | 81.8, 75.6, 88.7 | 83.0, 78.3, 87.2 | 82.6, 77.2, 85.6 | 90.0, 90.0, 90.0 |
| Resolution (Å) | 25.06-2.29 | 34.03-1.97 | 24.95-2.10 | 48.52-2.25 |
| *R*<sub>meas</sub> (%) | 0.133 (0.215) | 0.053 (0.334) | 0.145 (0.759) | 0.100 (0.656) |
| *I*/σ*I* | 7.61 (4.39) | 27.3 (4.36) | 7.52 (0.563) | 13.7 (0.846) |
| *cc*1/2 | (0.915) | (0.934) | (0.646) | (0.616) |
| Completeness (%) | 96.6 (97.3) | 99.7 (98.2) | 98.2 (91.2) | 98.5 (86.7) |
| Redundancy | 2.5 (2.6) | 3.9 (3.6) | 3.3 (1.8) | 5.6 (2.4) |
| **Refinement** |  |  |  |  |
| Resolution (Å) | 25.06-2.29 | 34.03-1.97 | 24.95-2.10 | 48.52-2.25 |
| No. reflections | 29791 | 76015 | 35000 | 63992 |
| *R*<sub>work</sub>/*R*<sub>free</sub> (%) | 22.96/25.08 | 18.52/20.33 | 25.34/27.44 | 21.49/24.79 |
| No. atoms |  |  |  |  |
| Protein | 4156 | 4307 | 4294 | 4388 |
| DNA | 839 | 836 | 836 | 1674 |
| Water | 274 | 515 | 298 | 252 |
| B-factors (Å<sup>2</sup>) |  |  |  |  |
| Protein | 35.79 | 27.84 | 34.39 | 29.26 |
| DNA | 52.26 | 45.55 | 52.53 | 59.12 |
| Water | 37.99 | 34.64 | 34.04 | 31.26 |
| R.m.s deviations |  |  |  |  |
| Bond length (Å) | 0.012 | 0.011 | 0.012 | 0.013 |
| Bond angles (°) | 1.705 | 1.390 | 1.754 | 1.816 |
| PDB ID | 9DP1 | 9DP2 | 9DP3 | 9DP4 |

We next crystallized APE1_N174A_ with the same AP DNA in the presence of MnCl_2_, promoting APE1 cleavage of the AP-site (31,42). From these crystals, we obtained a 1.97 Å structure of the APE1_N174A_ mutant product complex in the P1 space group (Table 2). Within the active site of the APE1_N174A_ product complex, there is clear density corresponding to the backbone being cleaved and Mn^2+^ bound (Fig. 3A-B). We overlayed the APE1_N174A_ mutant product complex with the APE1_WT_ product complex and observed similar active site conformations (RMSD=0.208 of 2449 atoms, Fig. 3C) (28). An exception was active site residue N212, which was positioned differently in the APE1_N174A_ mutant compared to APE1_WT_. Prior to cleavage in APE1_WT_, the carbonyl group of N212 orients the nucleophilic water through a hydrogen bonding interaction (Fig. 3D, 1B). Following cleavage, N212 undergoes a rotameric shift, positioning its amine group within hydrogen bonding distance of the 5’-phosphate terminus (Fig. 3D, 1C). In the APE1_N174A_ mutant, N212 fails to make this rotameric shift and remains in the same position as in the substrate structure (Fig. 3B, 3D). We expect that N174A mutation causes an unfavorable environment for the N212 rotameric shift, but that the N212 rotameric shift is not essential to cleavage.

**Figure 3.**
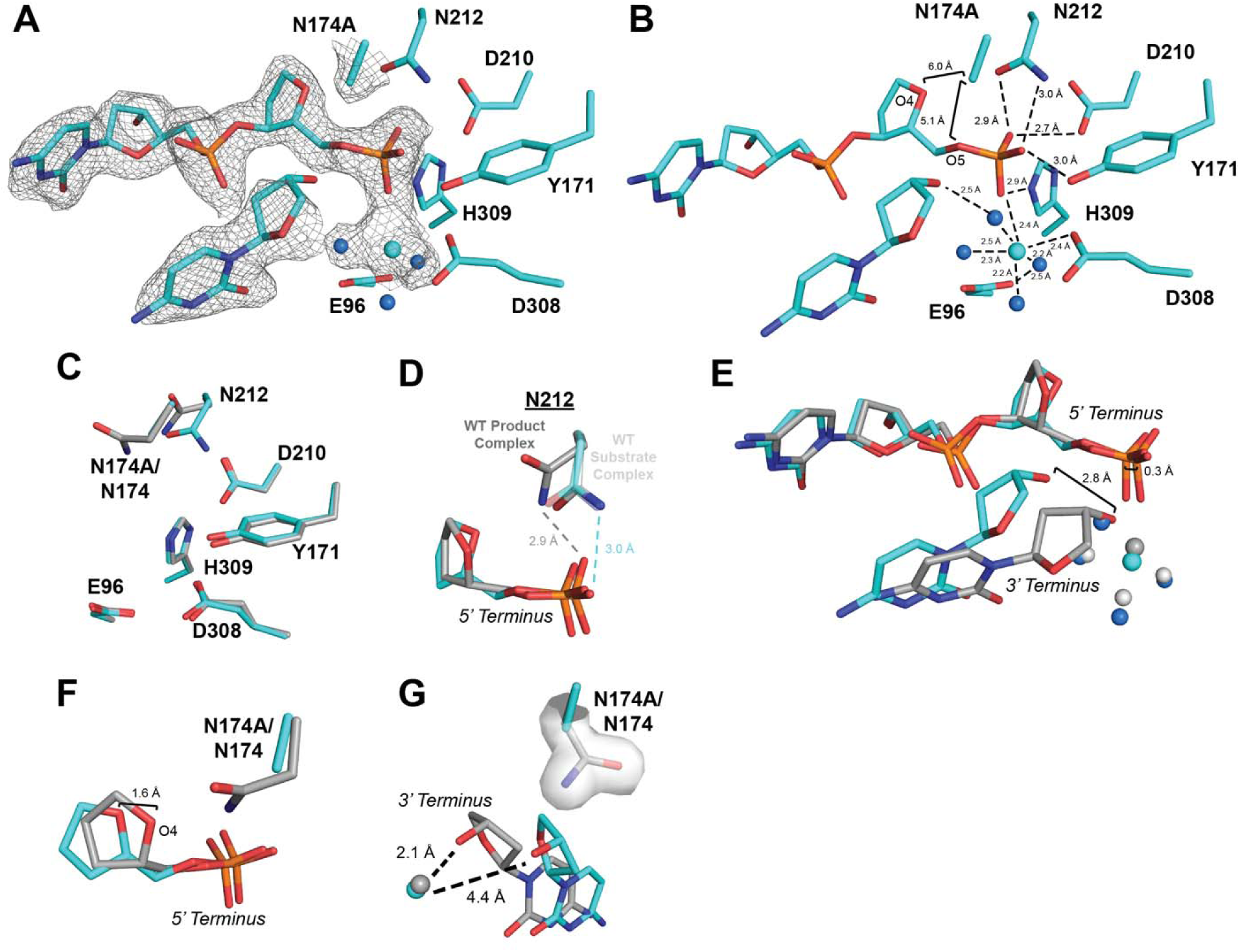
The APE1_N174A_ mutant product complex. (A-B) Active site of the APE1_N174A_ product complex crystal structure in cyan. Density is shown as grey mesh. Waters are shown as blue spheres and Mn^2+^ is shown as a cyan sphere. (C) Active site residues of the APE1_N174A_ product complex crystal structure in cyan compared to the active site of APE1_WT_ product complex shown in gray (PDB ID: 5DFF). (D) Position of N212 in the APE1_WT_ substrate complex crystal structure in light gray (PDB ID: 5DFI) and the APE1_WT_ product complex in dark grey (PDB ID: 5DFF). The rotameric position of N212 in the APE1_N174A_ product complex resembles the APE1_WT_ substrate complex. (E) Comparison of product DNA in the APE1_N174A_ product complex crystal structure in cyan compared to the APE1_WT_ product complex in gray. Mn^2+^ is shown as a cyan sphere and waters are blue spheres for APE1_N174A_; Mg^2+^ is shown as a dark grey sphere and waters are light grey spheres for APE1_WT_. (F) O4 of the AP-site shifts 1.6 Å away from N174A in the APE1_N174A_ mutant compared to APE1_WT_. (G) N174A mutation increases the distance between Mn ^2+^ ion and the 3’ hydroxyl terminus compared to APE1_WT_. Mn^2+^ in the APE1_N174A_ complex is shown as a cyan sphere and Mg^2+^ in the APE1_WT_ complex is shown as a dark grey sphere.

Other than N212, the remaining active site residues in the APE1_N174A_ mutant do not differ from APE1_WT_ (Fig 3C). In the APE1_N174A_ mutant product complex, however, we also observed changes in the positioning of the cleaved termini of the AP DNA (Fig. 3E). In APE1_WT_, N174 is within hydrogen bonding distance of the 5’ phosphate terminus at O4 and O5 (Fig. 1C). Hydrogen bonding is absent in the APE1_N174A_ mutant, resulting in a 1.6 Å shift of the 5’ phosphate terminus away from residue N174A compared to WT (Fig. 3F). We also observed a substantial shift of the 3’ hydroxyl terminus of the cleaved DNA in the APE1_N174A_ mutant (Fig. 3G). In APE1_WT_, N174 acts as a steric barrier that flanks the 3’ hydroxyl terminus, keeping it in position to coordinate with the metal cofactor (Fig. 3G). In the APE1_N174A_ mutant, residue 174 cannot act as a steric barrier due to its reduced side chain length. This allows the 3’ hydroxyl terminus to shift over 2 Å away from the active site metal ion (Fig. 3G). In APE1_WT_, the 3’ hydroxyl is 2.1 Å from its metal cofactor, but this distance increases to 4.4 Å in the APE1_N174A_ mutant (Fig. 3G). This finding indicates that in APE1_WT_, the side chain size of N174 acts as a steric barrier that, along with hydrogen bonding, properly positions the AP-site within the APE1 active site for cleavage.

### Negative Charge in the APE1_N174D_ Mutant Inhibits AP-site Cleavage

Of our N174 APE1 mutants, APE1_N174D_ exhibited the greatest decrease in the rate of AP-site cleavage. To determine which features of APE1_N174D_ contribute to this decrease, we sought to investigate the active site organization of the APE1_N174D_ mutant, both before and after cleavage. Unfortunately, we were not able to obtain APE1_N174D_ substrate crystals and only obtained crystals of a product complex (described below). To provide insight into the APE1_N174D_ substrate complex, we performed computational mutagenesis to generate an APE1_N174D_ substrate model using the high-resolution x-ray crystal structure of the APE1_WT_ substrate complex (42). Using this mutant model, we performed molecular dynamic simulations of the APE1_N174D_ substrate complex to investigate the conformation of N174D. Analysis of the simulations indicates that throughout the course of the simulation, the N174D rotamer fails to stabilize at a favored conformation in the APE1_N174D_ mutant active site (Fig. 4A) and is never within hydrogen bonding distance of the AP-site (Fig. 4B). This would be expected given the clash between the side chain oxygens of N174D and the highly electronegative DNA backbone. To compare the computationally generated APE1_N174D_ substrate complex to APE1_WT_, we also performed molecular dynamics simulations with the APE1_WT_ substrate complex. Analysis of the APE1_WT_ substrate complex simulations showed that unlike N174D, N174 of APE1_WT_ stabilizes at favored rotamer conformations (Fig. S5A) and was within hydrogen bonding distance of the AP-site for the entire simulation (Fig. S5B). This analysis confirms that our molecular dynamics simulations of the APE1_WT_ substrate complex is consistent with N174 hydrogen bonding to the phosphate backbone and demonstrates that the N174D mutation does not hydrogen bond to the phosphate backbone. This finding is consistent with our kinetics data, which indicated that the hydrogen bonding potential and charge of N174 in APE1_WT_ are necessary for catalysis.

**Figure 4.**
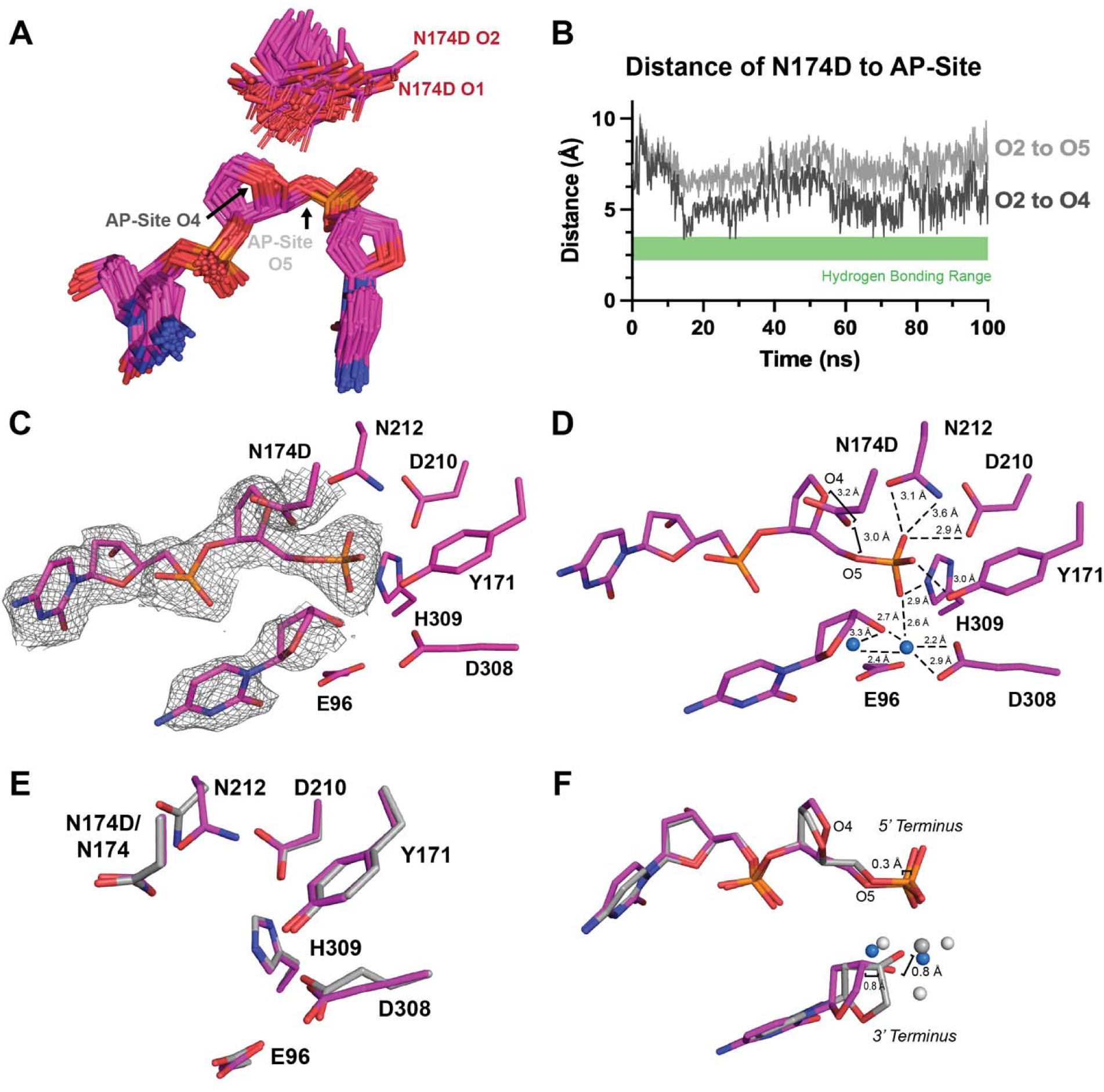
APE1_N174D_ substrate simulations and X-ray product complex. (A) Rotamer states of N174D in the APE1_N174D_ substrate complex determined by computational modeling. N174D rotamers fail to assume a primary rotameric state. (B) Distances determined between O2 of N174D and O4 and O5 of the AP-site in the APE1_N174D_ substrate complex determined by computational modeling were plotted to determine if N174D is within hydrogen bonding distance of the AP-site. Hydrogen bonding range is marked by the green box on distance plots as distances between 2.2 and 3.5 Å. (C-D) Active site of the APE1_N174D_ product complex crystal structure in magenta. Density is shown as grey mesh. Waters are shown as blue spheres. (E) Active site residues of the APE1_N174D_ product complex crystal structure in magenta compared to the active site of the APE1_WT_ product complex in gray (PDB ID: 5DFF). (F) The position of DNA termini within the APE1_N174D_ product complex crystal structure compared to the APE1_WT_ product complex in gray. Waters are blue spheres for APE1_N174D_; Mg^2+^ is shown as a dark grey sphere and waters are light grey for APE1_WT_.

Next, we crystallized the APE1_N174D_ mutant protein bound to a 21-mer double-stranded DNA oligo containing a centrally located AP-site analog tetrahydrofuran (THF). In these crystals, the AP-site has been cleaved by APE1 resulting in a product complex that diffracted to 2.10 Å resolution in the P1 space group (Table 2). Within this structure, both N174D and the AP-site are positioned in the active site with clear density corresponding to the backbone being cleaved, however there is no metal bound (Fig. 4C-D). We overlayed the APE1_N174D_ mutant product complex with the APE1_WT_ product complex and observed similar active site conformations (RMSD=0.335 of 2321 atoms, Fig. 4E) (28). Consistent with what we observed for the APE1_N174A_ mutant, residue N212 in the APE1_N174D_ mutant failed to undergo the rotameric shift observed in APE1_WT_ following product formation (Fig. 4E). The 5’ phosphate and 3’ hydroxyl DNA termini in the APE1_N174D_ mutant product complex are only moderately shifted compared to their positions in the APE1_WT_ product complex (Fig. 4F). These shifts are considerably smaller than those observed in the APE1_N174A_ mutant product complex, supporting the idea that the N174 side chain acts as a steric barrier to keep the cleaved 5’ and 3’ DNA termini within the APE1 active site.

Overall, the structure and simulations of our APE1_N174D_ mutant indicate that hydrogen bonding and side chain size of residue 174 stabilize DNA within the APE1 active site. As is the case for APE1_N174A_, we partially attribute the reduced cleavage rate of the APE1_N174D_ mutant to the lack of hydrogen bonding interactions observed in our simulations and structure. However, given that the rate of cleavage for the APE1_N174D_ mutant is even lower than in the APE1_N174A_ mutant, we hypothesize the charge of the N174D mutation may account for the remaining difference in cleavage rates between our mutants. To understand whether the charge of N174 mutation could impair cleavage, we visualized the averaged electrostatic potential of APE1_WT_ and each of our mutants prior to cleavage (Fig. 5). We found that a larger area of the APE1_N174D_ mutant active site surface was more negatively charged than APE1_WT._ This indicates that N174D mutation more broadly impacts the electrostatic environment of the APE1_N174D_ mutant active site and could impact catalysis. Therefore, this evidence from our mutants indicates that APE1 requires both hydrogen bonding potential and a suitable charge at residue 174 for cleavage.

**Figure 5.**
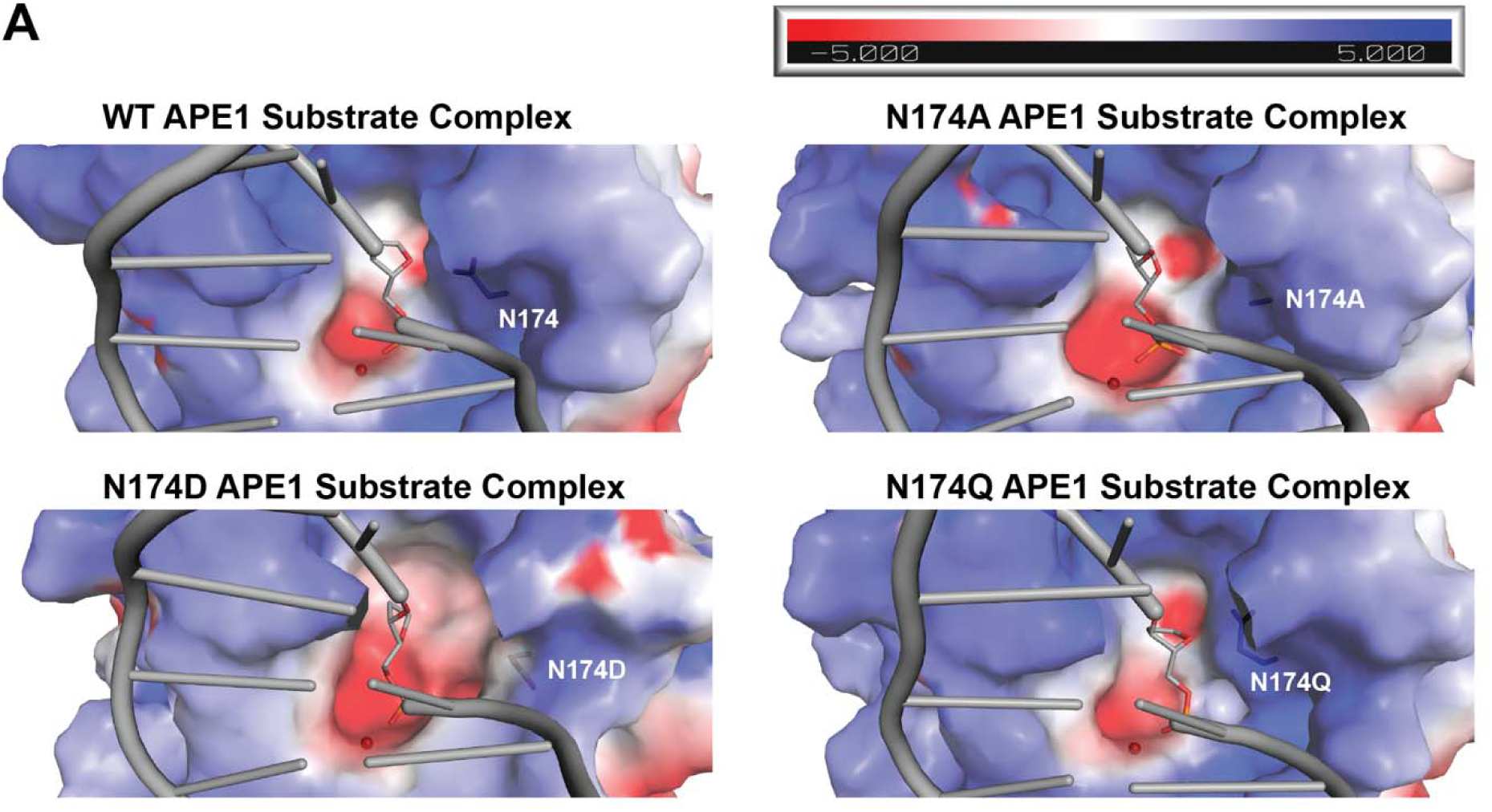
Effect of N174 mutation on the averaged electrostatic potential of the APE1 active site. The averaged electrostatic potential of APE1_WT_, APE1_N174A_, APE1_N174D_, and APE1_N174Q_ substrate complexes determined by computational simulations. Intercalating residues R177 and M270 of all models have been removed for clarity of the active site cavity. The active site of the APE1_N174D_ substrate complex is more negatively charged than that of APE1_WT_.

### Glutamine Stabilizes AP DNA in the APE1_N174Q_ Mutant and Mimics the WT Asparagine During AP-Site Cleavage

APE1_N174Q_ exhibited a decreased rate of AP-site cleavage, but to a lesser extent than either the APE1_N174A_ or APE1_N174D_. To understand why APE1_N174Q_ was able to partially restore cleavage activity, we attempted to investigate the active site organization of the APE1_N174Q_ mutant via x-ray crystallography. Unfortunately, we were not able to obtain suitable APE1_N174Q_ substrate complex crystals and only obtained crystals of a product complex (described below). Therefore, to provide insight into the APE1_N174Q_ substrate complex we performed computational mutagenesis to generate an APE1_N174Q_ substrate model using the high-resolution APE1_WT_ substrate complex (42). Analysis of molecular dynamics simulations of this complex indicates that N174Q primarily adopted rotameric conformations most similar to a single standard rotameric state, tp-100 (X_1_=180°, X_2_= 65°, X_3_= -100°, Fig. 6A) (43,44). Additionally, distances of the amine group of N174Q were within hydrogen bonding distance of the AP-site at O4 or O5 throughout the course of the simulation (Fig. 6B). This indicates the added side chain length of N174Q is accommodated in the APE1_N174Q_ active site, and that N174Q can still hydrogen bond to the DNA backbone. Therefore, the hydrogen bonding potential and charge at residue 174 are essential for proper catalysis in APE1.

**Figure 6.**
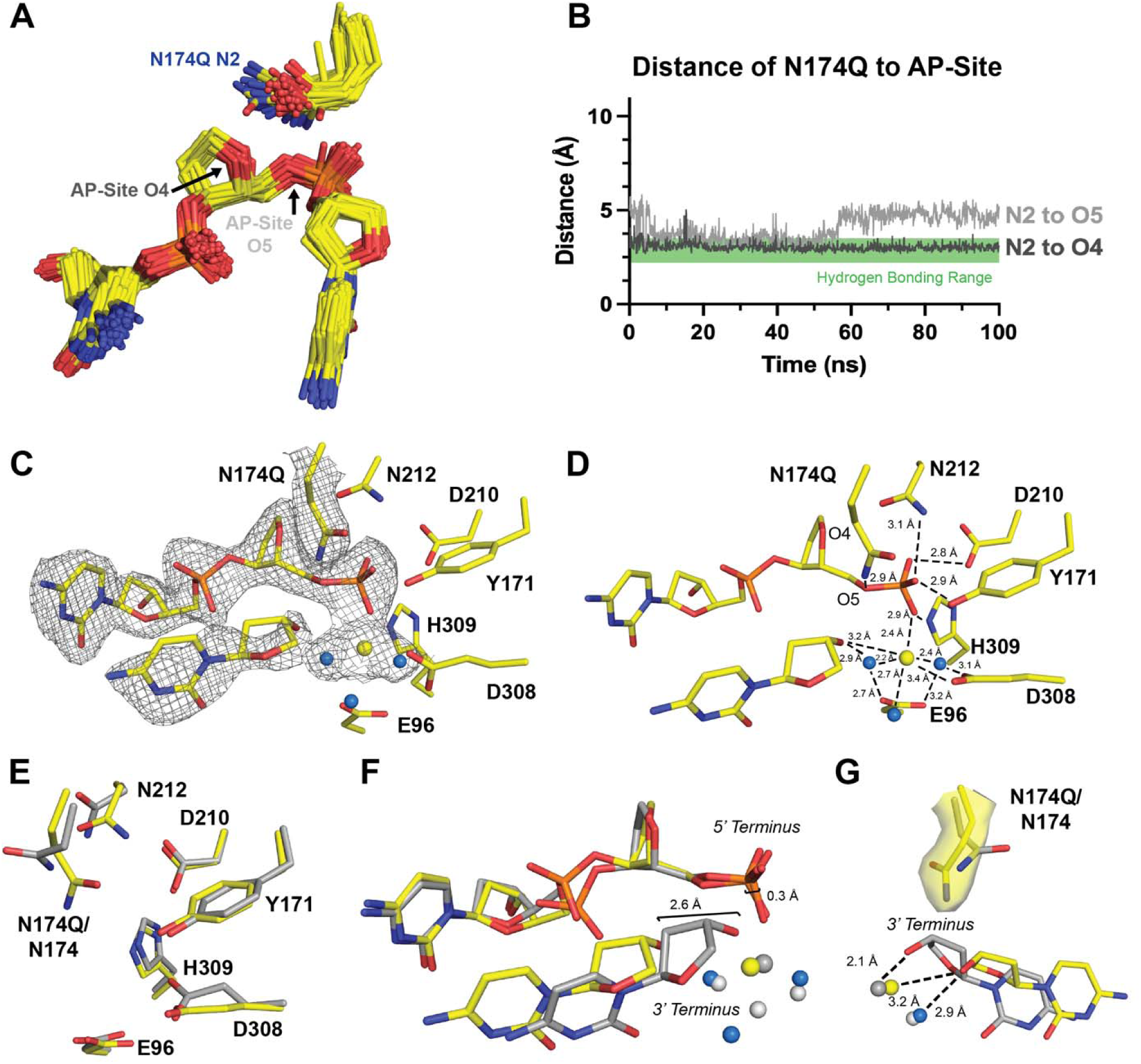
APE1_N174Q_ substrate simulations and X-ray product complex. (A) Rotamer states of N174Q in the APE1_N174Q_ substrate complex determined by computational modeling. Rotamers cluster near the standard rotamer conformation tp-100 (X_1_=180°, X_2_= 65°, X_3_= -100°). (B) Distances determined between N2 of N174Q and O4 and O5 of the AP-site in the APE1_N174Q_ substrate complex determined by computational modeling were plotted to determine if N174Q is within hydrogen bonding distance of the AP-site. Hydrogen bonding range is marked by the green box on distance plots as distances between 2.2 and 3.5 Å. (C-D) Active site of the APE1_N174Q_ product complex crystal structure in yellow. Density is shown as grey mesh. Waters are shown as blue spheres and Mg^2+^ is shown as a yellow sphere. (E) Active site residues of the APE1_N174Q_ product complex crystal structure in yellow compared to the active site of the APE1_WT_ product complex in gray (PDB ID: 5DFF). (F) DNA termini in the APE1_N174Q_ product complex in yellow compared to APE1_WT_ product complex in gray. Mg^2+^ is shown as a yellow sphere and waters are blue spheres for APE1_N174Q_; Mg^2+^ is shown as a dark grey sphere and waters are light grey spheres for APE1_WT_. (G) The longer side chain length of glutamine in the APE1_N174Q_ mutant compared to asparagine in APE1_WT_ displaces the 3’ hydroxyl terminus. Mg^2+^ is shown as a yellow sphere and waters are blue spheres for APE1_N174Q_; Mg^2+^ is shown as a dark grey sphere and waters are light grey spheres for APE1_WT_.

Finally, we crystallized the APE1_N174Q_ mutant bound to a 21-mer double-stranded DNA oligo containing a centrally located AP-site analog, THF. In these crystals, the backbone has been cleaved by APE1 resulting in a product complex that diffracted to 2.25 Å resolution in the P65 space group (Table 2) (31). The crystal form of the APE1_N174Q_ mutant product complex contains two nearly identical APE1_N174Q_:DNA complexes (RMSD=0.452 of 2329 atoms). We utilized the APE1_N174Q_:DNA complex in chain A for our structural analysis since it contained better electron density for the active site residues. The AP-site is positioned in the APE1_N174Q_ mutant active site with clear density corresponding to the backbone being cleaved and a Mg^2+^ ion bound (Fig. 6C-D). We overlayed the APE1_N174Q_ mutant product complex with the APE1_WT_ product complex and observed similar active site conformations (RMSD=0.510 of 2236 atoms, Fig. 6E) (28). The active site residues of APE1_N174Q_ align well with their positions in the APE1_WT_ active site (Fig. 6E). As in both the APE1_N174A_ and APE1_N174D_ mutant product complexes, residue N212 failed to undergo the rotameric shift observed in the APE1_WT_ product complex structure and remains in the substrate conformation (Fig. 6E). We did observe one additional change not present in either the APE1_N174A_ or APE1_N174D_ mutant product complexes at residue D308, which adopts a different rotameric confirmation than in APE1_WT_. This rotamer shift likely arises from minor changes in the position of the cleaved 3’ hydroxyl terminus and the coordinating metal ion (Fig. 6D). In the APE1_N174Q_ mutant, the 3’ hydroxyl terminus shifts 2.6 Å away from the mutant active site (Fig. 6F). Comparing APE1_WT_ to the APE1_N174Q_ mutant indicates that this shift in the 3’ hydroxyl terminus arises to allow the mutant active site to accommodate the extra side chain length of N174Q and prevent a clash between N174Q and the sugar moiety of the 3’ terminus (Fig. 6G).

Overall, the structure of our APE1_N174Q_ product complex is consistent with simulations of our APE1_N174Q_ substrate complex. Both the structure and simulations indicate that the hydrogen bonding potential and charge of N174Q assist the APE1_N174Q_ mutant in cleaving AP-sites. While the APE1_N174Q_ mutant retains better catalytic capabilities than other mutants, its activity is still less than APE1_WT_. We suggest that this difference likely results from a subtle displacement of the DNA within the APE1_N174Q_ mutant active site to accommodate the added side chain length of N174Q as observed in our product complex. Because the decrease in APE1_N174Q_ cleavage rate is moderate, N174Q may still be able to partially stabilize the APE1 cleavage reaction despite any potential DNA displacement. Therefore, in APE1_WT_ we expect the side chain length of N174 does contribute to cleavage, but that ultimately the hydrogen bonding potential and charge of N174 are most important for cleavage.

## Discussion

In this study, we demonstrate that the residue N174 is necessary for cleavage of AP-sites by APE1. Our study points to three key factors of residue N174 affecting cleavage of an AP-site: hydrogen bonding potential, charge, and size. Hydrogen bonding between N174 and the AP-site is required to stabilize AP DNA during APE1 cleavage. When hydrogen bonding is lost, as in our APE1_N174A_ and APE1_N174D_ mutants, we observed both decreased rates of cleavage and structural differences in the position of AP DNA (Table 1). Additionally, N174Q in our APE1_N174Q_ mutant was able to maintain similar hydrogen bonding to the AP-site as N174 in APE1_WT_ and the APE1_N174Q_ mutant had the least impaired cleavage rate of our N174 mutants (Table 1). Therefore, we conclude that hydrogen bonding between N174 and the AP-site is necessary for efficient cleavage of AP-sites by APE1.

Our study indicates that the hydrogen bonding potential between APE1_WT_ and AP DNA first observed in x-ray crystal structures (Fig. 1A-C) is indeed functionally relevant to APE1’s AP endonuclease activity. X-ray crystal structures of APE1 bound to other DNA substrates similarly indicate that N174 hydrogen bonds to DNA substrates during its exonuclease activity (45,46). This suggests to us that N174 could play an essential role not only during APE1 endonuclease cleavage of AP DNA but could also be implicated outside of AP-site processing such as during APE1’s exonuclease activities.

While hydrogen bonding explains how N174 stabilizes substrate DNA, we also expect the charge of N174 to contribute to the electrostatic environment of the APE1 active site enabling cleavage. Hydrogen bonding alone cannot account for differences observed in the cleavage rates of our mutants since neither N174A nor N174D are capable of hydrogen bonding to the AP-site. Yet our APE1_N174D_ mutant had a lower cleavage rate than our APE1_N174A_ mutant (Table 1). Visualization of the averaged electrostatic potential of our APE1 mutant active sites revealed that the APE1_N174D_ mutant active site was more negatively charged than any of the other APE1 mutants (Fig. 5). From this information, we hypothesize that additional negative charge within the APE1_N174D_ active site inhibits formation of the negatively charged phosphorane intermediate of the APE1 endonuclease reaction (Fig. 7A-B). Therefore, we suggest a model in which residue 174 in APE1 must contribute a favorable combination of charge and hydrogen bonding potential to promote formation of the phosphorane intermediate and allow the endonuclease reaction to proceed (Fig. S7B). We conclude that like hydrogen bonding, the charge of N174 is necessary for efficient cleavage of AP-sites by APE1 and likely other APE1 activities.

**Figure 7.**
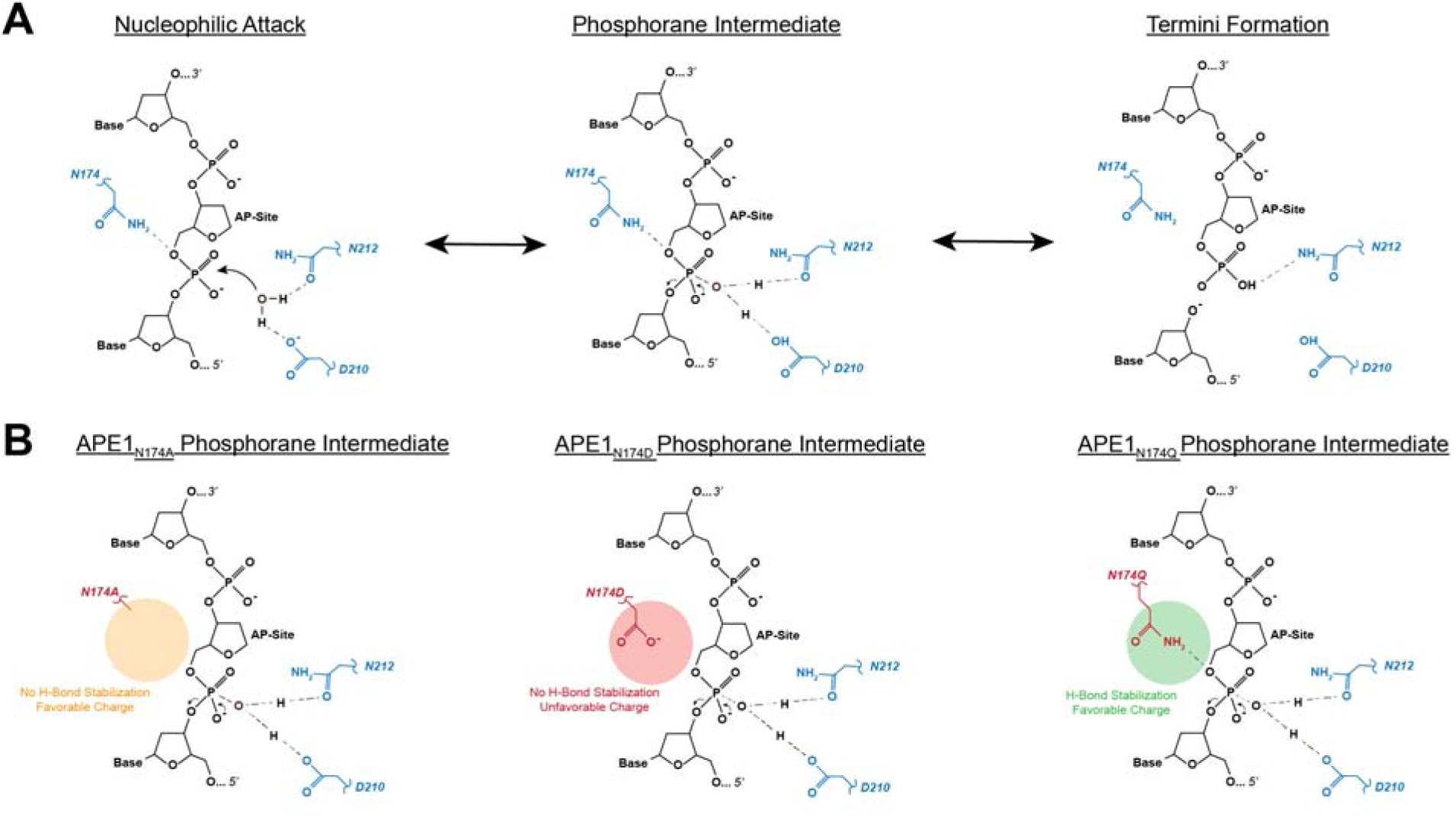
The effect of N174 and N174 mutation on the APE1 endonuclease reaction. (A) A simplified scheme* of the APE1 endonuclease reaction showing coordination of cleavage by D210 and N212 and stabilization of the reaction by N174. (B) Effect of N174 mutation on the phosphorane intermediate of the APE1 endonuclease reaction. *See the following references for a more complete scheme: (28,29)

Hydrogen bonding and charge are not the only evidence that N174 is essential to APE1 endonuclease activity and other potential repair functions. Analysis of our APE1 mutants, also indicates that the size of N174 in APE_WT_ contributes to APE1 cleavage of AP-sites. The smaller size of N174A led to considerable shifts in the position of the AP DNA within the APE1_N174A_ mutant active site, potentially impeding catalysis (Fig 2D, 3G). This suggests N174 in APE1_WT_ may behave as a steric barrier keeping substrate DNA within the APE1 active site during cleavage. However, the APE1_N174D_ mutant exhibited the lowest cleavage rate of all mutants compared to APE1_WT_ while maintaining side chain size (Table 1). Additionally, the APE1_N174Q_ mutant exhibited less impaired catalysis than the other mutants (Table 1), added the added side chain size was accommodated in the APE1_N174Q_ active site (Fig. 6). Cumulatively, this suggests that the size of N174 in APE1_WT_ is likely not as important for catalysis as hydrogen bonding or charge but does point to a moderate role of the side chain size in properly organizing the active site during catalysis. It also raises an interesting hypothesis of the N174 side chain size playing a larger role in other APE1 catalytic activities, such as the exonuclease activity.

As mentioned previously, N174 is highly conserved among the primary endonucleases from humans to zebrafish (SFig. 1). However, sequence alignment of human APE1 with the primary AP-endonuclease in *E. coli,* EXOIII (37–40), reveals alignment of N174 in APE1 with Q112 in EXOIII (SFig. 1). While EXOIII is the primary AP-endonuclease in *E. coli*, EXOIII is considered to have a more robust exonuclease than endonuclease activity (39,40). Inversely, APE1 is considered to have a more robust endonuclease activity when compared to its exonuclease activity (39,47). The active sites of APE1 and EXOIII are conserved with the exception of N174/Q112, implicating the difference in the size of N174 and Q112 in the variability of endonuclease vs. exonuclease activities across these enzymes. So far, differences in the exonuclease function between APE1 and EXOIII have been linked to bulky aromatic residues expected to stabilize the 3’ terminal base to be excised during exonuclease activity (48). But interestingly, Q112 does appear to affect EXOIII exonuclease activity on some substrates (49) suggesting that N174 could also affect the exonuclease activity of APE1 along with the suspected aromatic residues, which will be the focus of future studies.

In summary, analysis of N174 in APE1 indicates that AP-site cleavage by APE1 is dependent upon the hydrogen bonding, charge, and size of N174. We found that N174 stabilizes AP DNA during endonuclease cleavage via hydrogen bonding and side chain size, and that it promotes the endonuclease reaction via charge. Furthermore, we speculate that the size of N174 may be of unexplored interest concerning the differences in APE1’s endonuclease and exonuclease functions when compared to other AP-endonuclease homologs, in particular, EXOIII. These findings, taken together, provide novel characterization of an essential residue of the APE1 active site, and further our understanding of how APE1 coordinates its active site and DNA substrates for cleavage.

## Experimental Procedures

### DNA Substrates

All APE1 DNA substrates were prepared by annealing oligonucleotides purchased and purified by Integrated DNA Technologies (IDT). DNA substrates for kinetic analysis and electrophoretic mobility shift assays (EMSAs) were prepared by annealing a 5’ fluorescein amidite (FAM)-labeled oligonucleotide template containing a centrally located tetrahydrofuran-type (THF) AP-site with an unlabeled oligonucleotide complement (42). Oligonucleotides were annealed as follows: FAM-labeled and unlabeled oligonucleotides were mixed in water to a final concentration of 10 μM and 12 μM, respectively, heated to 95 **°**C, and cooled at a rate of 1 **°**C per minute until reaching 4 **°**C. Oligonucleotides for x-ray crystallography structure determination were annealed as follows: oligonucleotides were mixed in buffer containing 100 mM Tris, pH 7.5 and 20 mM MgCl_2_ to a final concentration of 2 mM, heated to 95 **°**C, and cooled at a rate of 1 **°**C per minute until reaching 4 **°**C. The 30-bp oligonucleotide sequences used for kinetic analysis and EMSAs were 5’*-CGT-TCG-CTG-ATG-CGC-XCG-ACG-GAT-CCG-CAT-3’ and 5’-ATG-CGG-ATC-CGT-CGA-GCG-CAT-CAG-CGA-ACG-3’ (where * denotes FAM-label and X denotes the THF AP-site) (42). The 21-bp oligonucleotide sequences used for x-ray crystallography structure determination for product structures were 5’-GGA-TCC-GTC-GGG-CGC-ATC-AGC-3’ and 5’-GCT-GAT-GCG-CXC-GAC-GGA-TCC-3’ (42).

### Protein Expression and Purification

Human full-length WT and N174 mutant APE1 was expressed and purified as previously described (45). Briefly, APE1 was expressed in NEB PLysY *E. coli* using an optimized PET28a vector. Cells were lysed via sonication and lysate was filtered with a 0.45 μm filter then purified using an ATKA-Pure FPLC with the following columns: Cytiva 5 mL Heparin HP affinity column, Cytiva 5 mL Resource S cation exchange column, and Cytiva HiPrep 16/60 Sephacryl S-200 HR gel filtration column. Proteins were concentrated to stock concentrations of approximately 5 mg/mL for kinetics and EMSA experiments and 20 mg/mL for x-ray crystallography experiments. Final protein concentration was determined using absorbance at 280 nm. Full-length WT and N174 mutant APE1 proteins were utilized for kinetic experiments and EMSAs. Truncated (Δ1-42)/C138A variants, which have been used previously and shown not to alter APE1 function, were utilized for crystallography studies (27,28,31).

### Single and Multiple Turnover Kinetic Analysis

Cleaved DNA formed by full length WT and N174 mutant APE1 formed over time was measured under both single turnover and multiple turnover conditions. Product formation was measured by visualizing FAM-labeled substrate on a 21% denaturing National Diagnostics Ultra-Pure Sequagel UreaGel gel. Single turnover conditions were utilized to approximate the cleavage rate constant, k_obs_. Multiple turnover conditions were utilized to approximate the cleavage rate constant, *k_obs_*, and steady state rate constant, *k_ss_*. Cleavage reactions were initiated by mixing substrate DNA with APE1 and allowing the reaction to proceed until quenching at pre-determined timepoints. Single-turnover reactions occurred under the following conditions: 50 mM HEPES, pH 7.5, 100 mM KCl, 5 mM MgCl_2_, 0.1 mg/mL BSA, 50 nM DNA, and 500 nM APE1. Multiple-turnover reactions occurred under the following conditions: 50 mM HEPES, pH 7.5, 100 mM KCl, 5 mM MgCl_2_, 0.1 mg/mL BSA, 100 nM DNA, and 30 nM APE1. Reactions were quenched with 300 mM EDTA, pH 8.0 on a Kintec Rapid Quench Flow, or with formamide loading dye containing 78% w/v formamide, 100 mM EDTA, 0.5 mg/mL bromophenol blue, and 0.25 mg/mL xylene cyanol. Quenched reactions were incubated with formamide loading dye at 95 °C for 5 minutes then run on a 21% denaturing National Diagnostics Ultra-Pure Sequagel UreaGel gel to resolve the fraction of cleaved DNA product from substrate, and visualized using a Typhoon imager. Product and substrate bands in the gel were quantified by densitometry with ImageQuantTL (GE, v 8.1) or ImageJ to calculate the amount of product formation over time (50). The resulting curve was fit to Eq 1 for single-turnover reactions or to Eq. 2 for multiple-turnover reactions using Prism v9.1.

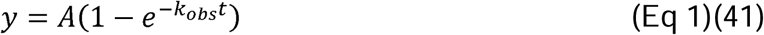

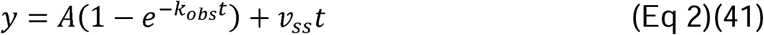

In Eq 1, *A* corresponds to the active fraction of enzyme and equals to the amplitude of product formation. In Eq 2, *A* corresponds to the active fraction of enzyme and equals the amplitude of product formation during the burst phase, and *v_ss_*is the steady state velocity where *k_ss_=v_ss_/A*.

As reported previously, the *k_obs_* of APE1_WT_ under single turnover conditions exceeds quantification using rapid quench flow instrumentation (41). Using this instrumentation, quenching APE1_WT_ cleavage of AP DNA under single turnover conditions after 0.0026 s resulted in approximately greater than 45% product. Therefore, we chose to present the *k_obs_* of APE1_WT_ under single turnover conditions as the previously determined minimum estimated rate constant necessary to exceed quenching under these conditions of at least 850 s^−1^ (41).

### Electrophoretic Mobility Shift Assay (EMSA)

WT and mutant APE1 affinity for a 30 bp DNA substrate containing an AP-site was determined by mixing APE1 with 2 nM DNA and allowing samples to incubate for at least 20 minutes to approach equilibrium. Concentrations of APE1 varied between 0-100 nM for APE1_WT_ and the APE1_N174Q_ mutant, and between 0-300 nM for the APE1_N174A_ and APE1_N174D_ mutants. Experiments were conducted under the following conditions: 50 mM Tris, pH 8.0, 1 mM EDTA, 0.2 mg/mL BSA, 50 mg/mL sucrose, 0.5 mg/mL bromophenol blue, and 1 mM DTT. Samples were run on a 10% 59:1 polyacrylamide native gel to determine APE1 bound and unbound fractions of DNA. FAM-labeled DNA was visualized using a Typhoon Imager. Bound and unbound fractions were quantified by densitometry with ImageQuant TL (GE, v 8.1). The resulting curve of bound DNA determined by ligand depletion vs. APE1 concentration was fit to Eq 3 using Prism v9.1 to determine the apparent binding affinity, *K*_D App_ (41).

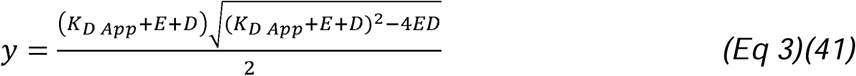

In Eq 3, *E* corresponds to the concentration of APE1 and *D* corresponds to the concentration of DNA.

### X-Ray Crystallography Structure Determination

APE1 structures were crystallized utilizing constructs with a 42 amino acid N-terminal truncation, a C138A mutation, and N174A, D, or Q mutations (31). To crystallize the APE1_N174A_ substrate complex, 0.56 nM DNA containing an AP-site was mixed with 16 mg/mL APE1_N174A_. Crystals were generated via sitting drop vapor diffusion using 2 μL of the above protein/DNA mix combined with 2 μL reservoir solution (100 mM Sodium Citrate, pH 5.0, 200 mM MgCl_2_, 16%-21% PEG 20,000). Resultant crystals were transferred to a cryoprotectant solution containing 75% reservoir solution supplemented with 20% ethylene glycol and 5% MgCl_2_, flash frozen, and subjected to x-ray diffraction. To crystalize the APE1_N174A_ product complex, the protein/DNA mix was first incubated with 50 mM MnCl_2_ for 100 mins and then crystalized as above. Resultant crystals were transferred to a cryoprotectant solution containing 75% reservoir solution supplemented with 20% ethylene glycol and 5% MnCl_2_, flash frozen, and subjected to x-ray diffraction. To crystallize the APE1_N174D_ product complex, 0.56 nM DNA containing an AP-site was mixed with 10 mg/mL APE1_N174D_. Crystals were generated via sitting drop vapor diffusion using 2 μL of the above protein/DNA mix combined with 2 μL reservoir solution (200 mM Lithium Sulfate and 15%-25% PEG 3350). Resultant crystals were transferred to a cryoprotectant solution containing 75% reservoir solution supplemented with 20% ethylene glycol and 5% MgCl_2_, flash frozen, and subjected to x-ray diffraction. To crystallize the APE1_N174Q_ product complex, 0.56 nM DNA containing an AP-site was mixed with 10 mg/mL APE1_N174Q_. Crystals were generated via sitting drop vapor diffusion using 2 μL of the above protein/DNA mix combined with 2 μL reservoir solution (100 mM HEPES Free Acid, 200 mM Ammonium Acetate, and 25% PEG 3350). Resultant crystals were transferred to a cryoprotectant solution containing 75% reservoir solution supplemented with 20% ethylene glycol and 5% MgCl_2_, flash frozen, and subjected to x-ray diffraction. APE1_N174D_ and APE1_N174Q_ with a phosphorothioate-containing AP substrate resisted crystallization. The APE1_N174A_ substrate structure was collected at the Lawrence Berkley National Laboratory Advanced Light Source using Beamline 4.2.2. The APE1_N174A_, APE1_N174D_, and APE1_N174Q_ product complex structures were collected at 100 K on a Rigaku MicroMax-007 HF rotating anode diffractometer system at a wavelength of 1.54 Å utilizing a Dectris Pilatus3R 200K-A detector. The software HKL3000 (v 705c, HKL Research Inc.) was used to process and scale diffraction data following collection from all x-ray diffractions sources. Initial models were determined using molecular replacement in PHENIX (v 1.19.2-4158-000) using a previously determined APE1:DNA product complex structure (PDB: 5DFF). Refinement and model building were done with PHENIX and Coot (v 0.9), respectively, and figures were made using PyMOL (v 2.5.1, Schrödinger LLC) (51,52).

### Computational Mutagenesis and Simulations

Computational mutagenesis was performed using *leap* modular of AMBER v16 suite of programs for biomolecular simulations (53). The side chain was deleted in the input file provided to the *leap* module, which generated the side chain conformation for mutant residue based on internal template. Molecular dynamics (MD) simulations were performed for APE1:AP DNA complexes in explicit water solvent. Computational modeling was performed to determine rotamer states of residue 174 and resulting distance to the AP-site in the APE1_WT_, APE1_N174D_, and APE1_N174Q_ substrate complexes, using an approach similar to those described in previous studies (54,55). Modeling of the APE1_N174A_ complex was not performed since alanine is unable to assume more than a single rotamer state. Model preparation and simulations were performed using the AMBER v16 suite of programs for biomolecular simulations (53). AMBER’s *ff14SB* (56) force-fields were used for all simulations. MD simulations were performed using NVIDIA graphical processing units (GPUs) and AMBER’s *pmemd.cuda* simulation engine using the Agarwal lab’s previously published protocols (57,58).

A total of 3 separate simulations were performed for APE1_WT_, APE1_N174D_ and APE1_N174Q_ based on the X-ray crystal structure of APE1_WT_ in complex with substrate DNA (PDB ID: 5DGI). The DNA sequence and APE1 constructs used in this study were similar to the study that produced the referenced structure (28). Any missing hydrogen atoms were added by AMBER’s *tleap* program. The N174D and N174Q mutations were created by AMBER’s *tleap* program. After processing the coordinates of the protein and substrate, all systems were neutralized by addition of counter-ions and the resulting system was solvated in a rectangular box of SPC/E water with a 10 Å minimum distance between the protein and the edge of the periodic box. The prepared systems were equilibrated using a protocol described previously (59). The equilibrated systems were then used to run 100 nanoseconds (ns) of production MD under constant energy conditions (NVE ensemble). The use of NVE ensemble is preferred as it offers better computational stability and performance (60). The production simulations were performed at a temperature of 300 K. As NVE ensemble was used for production runs, these values correspond to the initial temperature at the start of simulations. A temperature adjusting thermostat was not used in simulations. Over the course of the 100 ns simulation, the temperature fluctuated ± 5 K around the 300 K target, which is typical for well equilibrated systems.

As the goal of the computer simulations was to investigate the rotameric states of the protein residues, the DNA structure was restrained using positional restraints (1 kcal/mol/Å^2^ on all the DNA atoms). This allowed minimal movement in the DNA structure, while allowing protein residues to explore different possible conformations. A total of 1,000 conformational snapshots (stored every 100 picoseconds) collected for each system was used for analysis. The conformational snapshots were used for the rotameric sidechain evaluations and electrostatic potential calculations.

### Electrostatic Potential Calculations

For the 1,000 structures extracted above the electrostatic surfaces were computed using the Adaptive Poisson-Boltzmann Solver (APBS) software (61). The surfaces for the 1,000 structures were averaged to correspond to the representative electrostatic surface for the entire MD simulation trajectory.

### Multiple Sequence Alignment

Full amino acid sequences for human APE1 (Uniprot ID: P27695), chimpanzee APE1 (Uniprot: A2T6Y4), mouse APE1 (Uniprot: P28352), zebrafish APE1 (Uniprot: A0MTA1), and *E. coli* Exonuclease III (P09030) were obtained from UniProtKB/Swiss-Prot and aligned using EMBL-EBI ClustalOmega (CLUSTAL O(1.2.4)) (62,63).

Alignment output was formatted as ClustalW with character counts using default options for this output format. The alignment generated by ClustalOmega was visualized for publication using JalView (v 2.11.4.1) (64) and colored by consensus sequence where sequence identity was greater than or equal to 60% across the five species.

## Supporting information

Supporting Information, contains Figures S1-S5

## Data Availability

Additional information on X-ray crystal structure determination and structure coordinates is available from the Protein Data Bank (PDB codes: 9DP1, 9DP2, 9DP3, and 9DP4).

## Acknowledgements

Beamline 4.2.2 of the Advanced Light Source, a DOE Office of Science User Facility under Contract No. DE-AC02-05CH11231, is supported in part by the ALS-ENABLE program funded by the National Institutes of Health, National Institute of General Medical Sciences, grant P30 GM124169-01. Special thanks to Beamline Director Jay Nix. N.M.H. is a Howard Hughes Medical Institute Fellow of the Damon Runyon Cancer Research Foundation, DRG-2499-23.

## Funding Information

This work was supported by grants from the National Institute of General Medical Sciences [R35GM128562 to B.D.F. and R01GM148886 to P.K.A.]. The content is solely the responsibility of the authors and does not necessarily represent the official views of the National Institutes of Health.

## Notes

https://www.rcsb.org/structure/9DP1

https://www.rcsb.org/structure/9DP2

https://www.rcsb.org/structure/9DP3

https://www.rcsb.org/structure/9DP4

### Competing Interest Statement

The authors have declared no competing interest.

