## Supporting Information, contains Figures S1-S5 for "APE1 active site residue Asn174 stabilizes the AP-site and is essential for catalysis"

Supporting Information Includes:

Supplemental Figures S1-S5

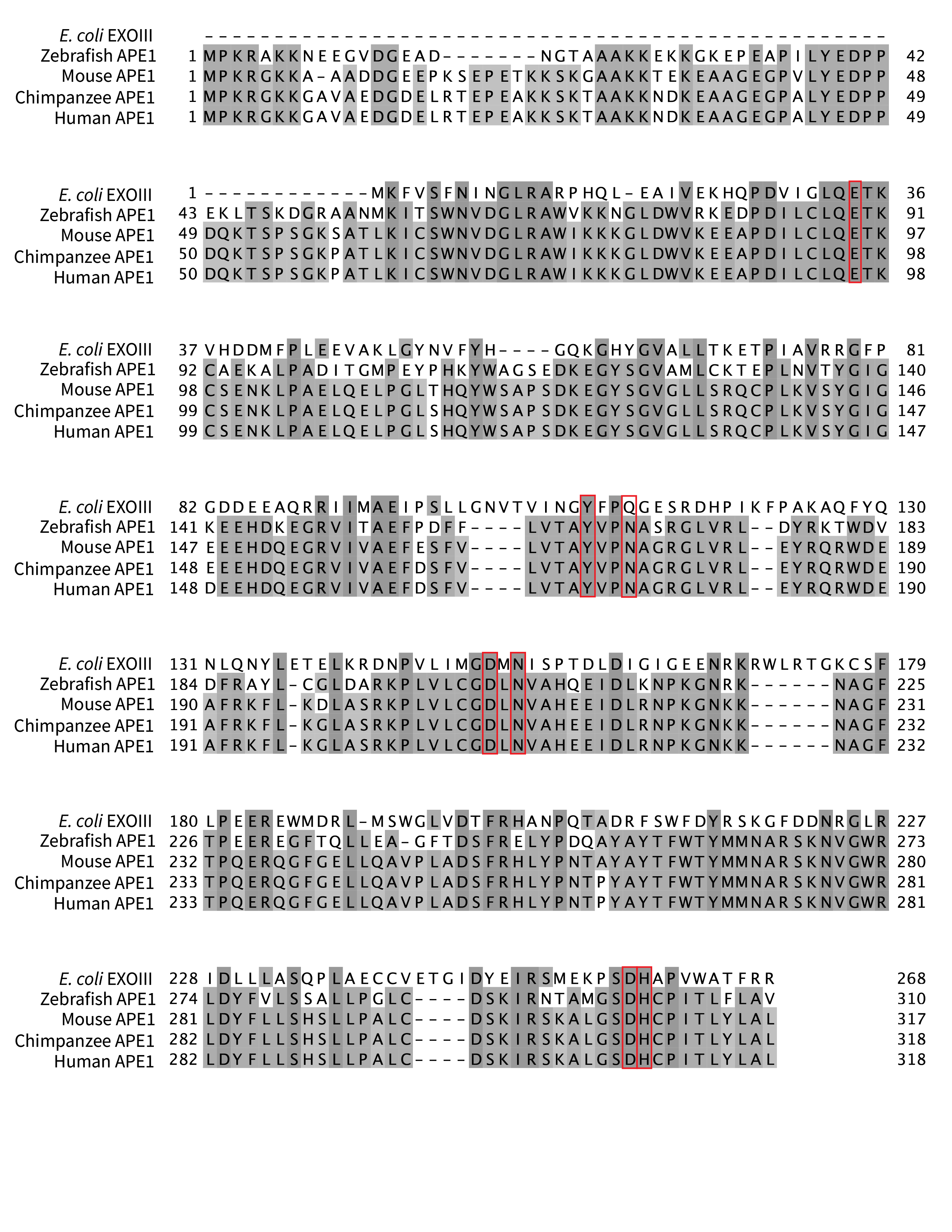

Supplemental Figure S1: **Multiple sequence alignment of primary AP-endonucleases across five selected species**

Full amino acid sequences for human APE1 (Uniprot ID: P27695), chimpanzee APE1 (Uniprot: A2T6Y4), mouse APE1 (Uniprot: P28352), zebrafish APE1 (Uniprot: A0MTA1), and E. coli Exonuclease III (P09030) were aligned using ClustalOmega. The multiple sequence alignment was visualized using JalView v 2.11.4.1 and colored by consensus sequence where sequence identity was greater than or equal to 60% across the five species. Red boxes outline active site residues.

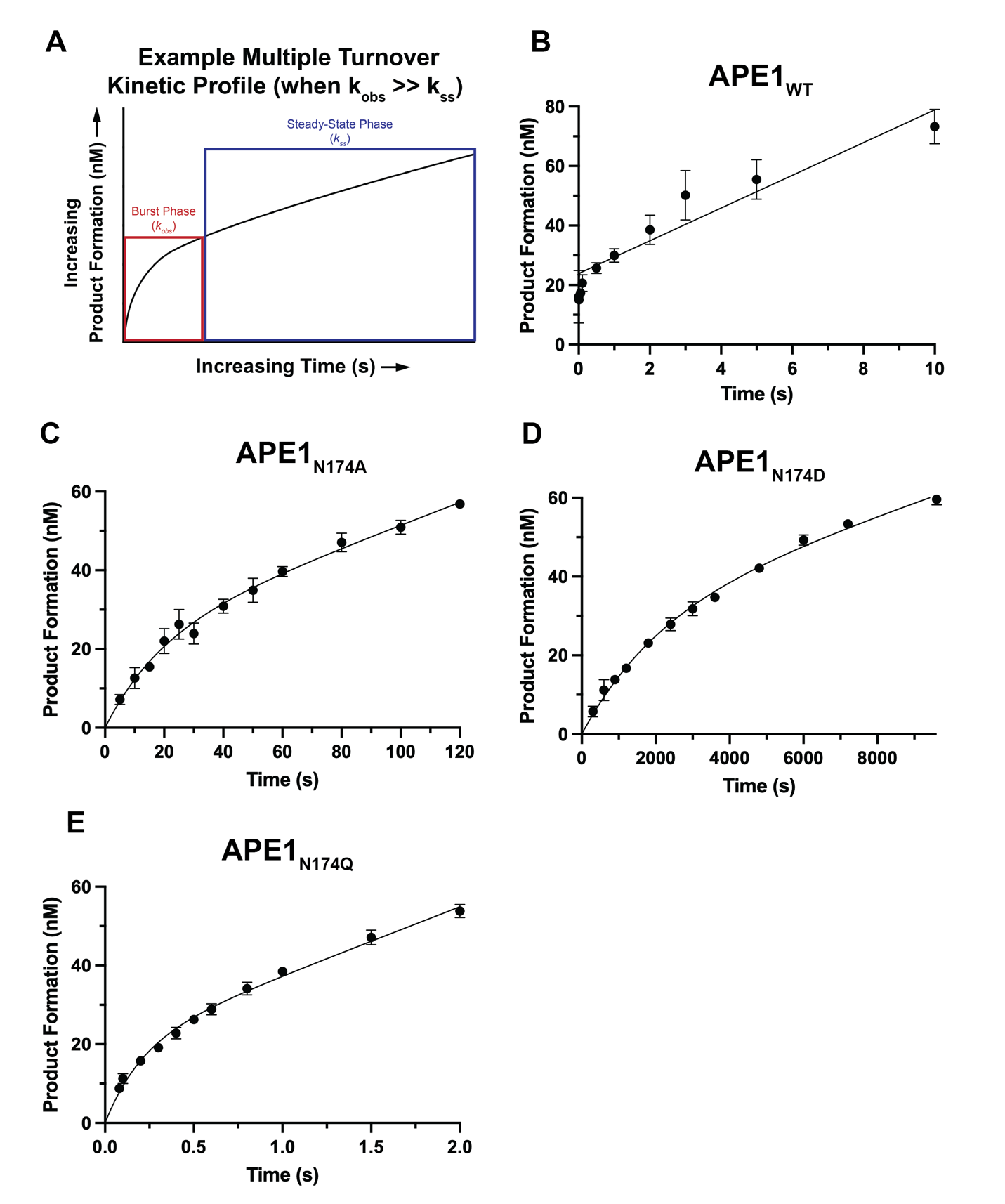

Supplemental Figure S2: **Fits of multiple turnover analysis of APE1 N174 mutants**

The amount of product formation (mean ± SE, N=3) over time under multiple turnover conditions was analyzed to determine the cleavage rate constant, *k_obs_*, and steady-state rate constant, *k_ss_*, for APE1_WT_, APE1_N174A_, APE1_N174D_, and APE1_N174Q_. Error bars are smaller than can be displayed when not present. (A) The cleavage rate constant, *k_obs_*, corresponds to the burst phase of product formation, and is followed by the steady state phase, corresponding to product release and the steady-state rate constant, k_ss_. The biphasic fits of multiple turnover kinetic analysis are shown for (B) APE1_WT_, (C) APE1_N174A_, (D) APE1_N174D_, and (E) APE1_N174Q_.

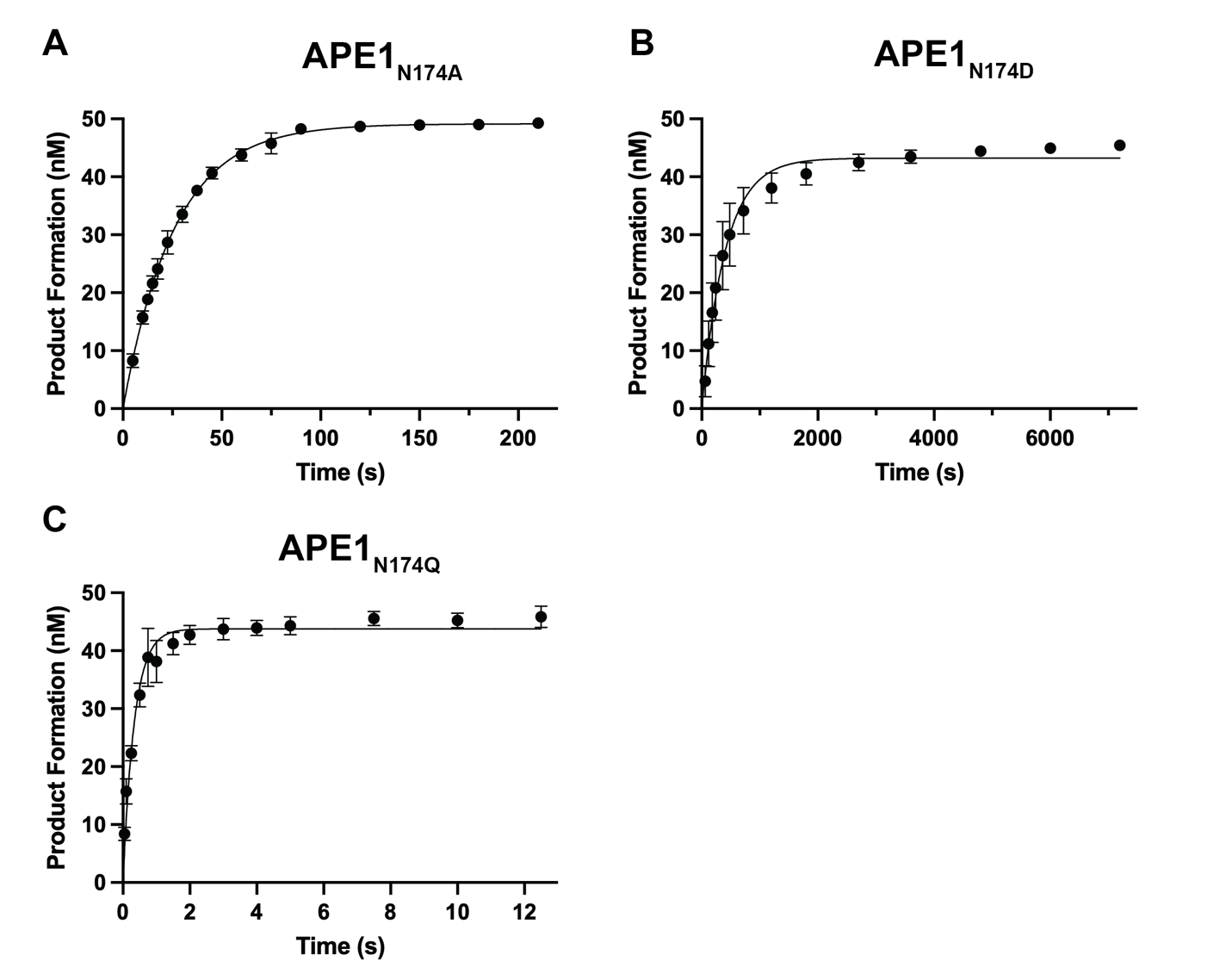

Supplemental Figure S3: **Fits of single turnover analysis of APE1 N174 mutants**

The amount of product formation (mean ± SE, N=3) over time under single turnover conditions was analyzed to determine the cleavage rate constant, *k_obs_*, for APE1 N174 mutants. Error bars are smaller than can be displayed when not present. The fits of single turnover kinetic analysis for (A) the APE1_N174A_ mutant, (B) the APE1_N174D_ mutant, and (C) the APE1_N174Q_ mutant are shown above.

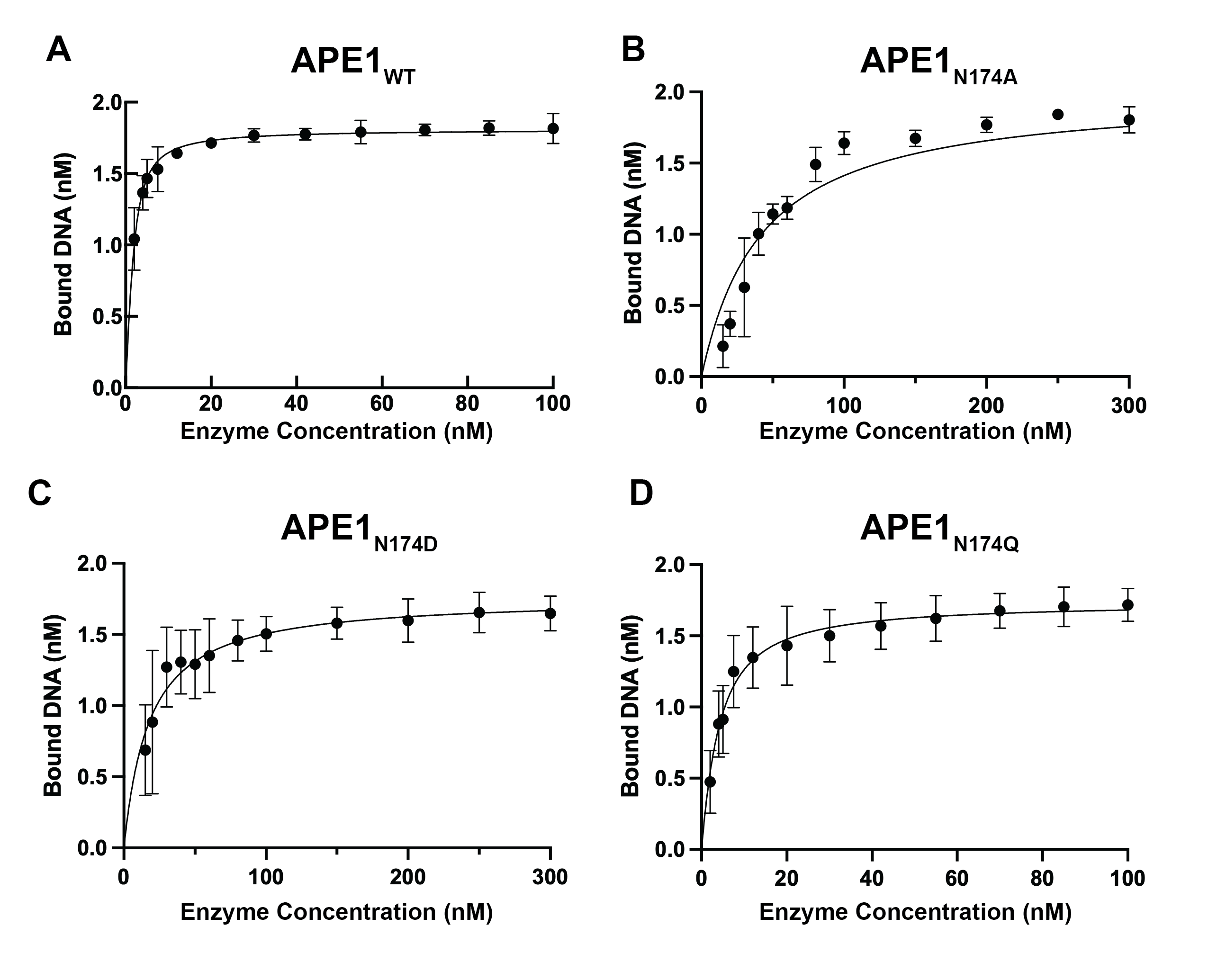

Supplemental Figure S4: **Fits of electrophoretic mobility shift assays (EMSAs) of APE1 N174 mutants**

The amount of DNA bound (mean ± SE, N=3) by APE1 mutants vs the concentration of mutant APE1 was analyzed to determine the apparent binding affinity, K_D App_, for APE1 and all APE1 N174 mutants. Error bars are smaller than can be displayed when not present. The fits of apparent binding affinity analysis for (A) APE1_WT_, (B) the APE1_N174A_ mutant, (C) the APE1_N174D_ mutant, and (D) the APE1_N174Q_ mutant are shown above.

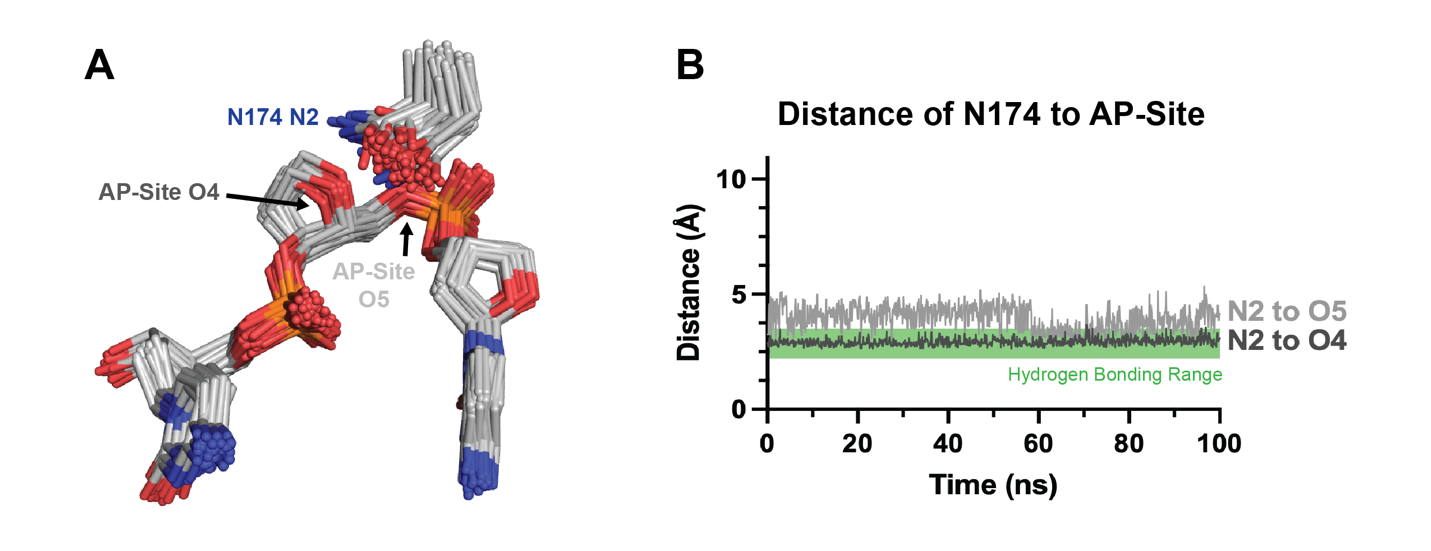

Supplemental Figure S5: **Analysis of APE1_WT_ rotamer conformation and hydrogen bonding distances from computational simulations**

Computational modeling was performed to determine rotamer states of residue 174 and the resulting distances to the AP-site in the APE1_WT_ substrate complex. (A) Rotamer states of N174 in the APE1_WT_ substrate complex determined by computational modeling. Rotamers cluster near the standard rotamer states m-20 and m-80 (both Χ_1_=-71). (B) The distances between N2 of N174 and O4 and O5 of the AP-site were plotted to determine if N174 is within hydrogen bonding distance of the AP-site in the APE1_WT_ substrate complex. Hydrogen bonding range is marked by the green box on distance plots as distances between 2.2 and 3.5 Å. Here, N2 of N174 in the APE1_WT_ substrate complex enters hydrogen bonding distance of either O4 or O5 of the AP-site in our computational simulations.
